# Biodiversity monitoring in Europe: user and policy needs

**DOI:** 10.1101/2023.07.12.548673

**Authors:** Hannah Moersberger, Jose Valdez, Juliette G. C. Martin, Jessica Junker, Ivelina Georgieva, Silke Bauer, Pedro Beja, Tom D. Breeze, Miguel Fernandez, Néstor Fernández, Lluís Brotons, Ute Jandt, Helge Bruelheide, W. Daniel Kissling, Christian Langer, Camino Liquete, Maria Lumbierres, Anne Lyche Solheim, Joachim Maes, Alejandra Morán-Ordóñez, Francisco Moreira, Guy Pe’er, Joana Santana, Judy Shamoun-Baranes, Bruno Smets, César Capinha, Ian McCallum, Henrique M. Pereira, Aletta Bonn

## Abstract

To implement the goals of the 2030 Global Biodiversity Framework, the European Biodiversity Strategy and the EU Green Deal, biodiversity monitoring is a pivotal instrument to achieve accountability and progress in conservation. Monitoring efforts in Europe, however, suffer from gaps and biases in taxonomy, spatial coverage, and temporal resolution, resulting in fragmented and disconnected data which does not provide sufficient evidence for policy making. To assess user and policy needs in biodiversity monitoring, we employed with EuropaBON a four-step user-centred stakeholder engagement process, including an international public stakeholder workshop, a standardised online survey, semi-structured interviews, and an expert meeting with representatives of EU member states, the European Commission and the European Environment Agency. The resulting insights into national and European biodiversity monitoring schemes identify policy needs, current challenges and potential solutions. Based on this in-depth policy and science stakeholder assessment, we recommend the establishment of a European Biodiversity Observation Network through a permanent Biodiversity Monitoring and Coordinating Centre to optimise existing observation efforts, harmonise data, and enhance our ability to predict and respond to key challenges related to biodiversity loss in a changing climate in Europe.

## 1. Introduction

Biodiversity monitoring forms a critical component of successful conservation policy and management. Successfully implementing, assessing and evaluating policy effectiveness and management relies on the availability of high-quality, scientifically robust, and reliable monitoring data, as well as their underlying methods for collection and analysis (Perino et al. 2022). Country borders, however, combined with sociopolitical barriers (such as language or mistrust between sectors) result in disconnected monitoring efforts and incoherence of data (Kühl et al. 2020). This is a key hurdle for gaining a comprehensive understanding of the state and trends of biodiversity (Proença et al. 2017) and for informing as well as evaluating effective policy and management interventions. Therefore, biodiversity monitoring requires collaborative, integrative and multinational approaches.

The importance of effective biodiversity monitoring has been increasingly recognised by policy- and decision-makers. The recent adoption of the Global Biodiversity Framework (GBF) at the 15th Conference of the Parties (COP) of the Convention on Biological Diversity (CBD) represents a significant step forward in the global conservation and restoration of biodiversity. For the first time, a global monitoring framework has been proposed to more efficiently implement and measure progress towards global biodiversity goals and targets (CBD 2022). Similarly, the European Biodiversity Strategy 2030 places a key commitment to monitoring protected areas, and the European Commission has also announced its ambition to establish workflows for monitoring and reporting biodiversity trends as part of the new European biodiversity governance framework (Directorate-General for Environment 2021).

One of Europe’s most significant policy goals is the EU Biodiversity Strategy’s attempt to halt the decline of biodiversity and promote its recovery by 2030 (Directorate-General for Environment 2022). One way to achieve this is through the restoration of a significant portion of degraded ecosystems in order to provide long-term ecosystem services, for which the proposed EU’s new Nature Restoration Law could pave the way through legally-binding restoration targets (Directorate-General for Environment 2022). Thus, the various European biodiversity monitoring initiatives need to provide solid and integrated empirical evidence for the achievement of these policy goals and evaluate their effectiveness and impact over space and time. Determining the effectiveness and impact of biodiversity-related policies is also crucial to evaluating the success of conservation and restoration efforts in accordance with the EU Nature Directives, the Water Framework Directive (WFD), the Common Agricultural Policy (CAP), the targets set by the EU Biodiversity Strategy, and other cornerstones of EU legislation e.g. for climate and soil.

To fulfil these ambitions, the EU needs robust, reliable streams of biodiversity data across spatial and temporal scales. Nevertheless, current monitoring efforts in Europe suffer from limitations such as taxonomic, spatial, and temporal gaps and biases. They are often fragmented across ecosystems and habitats, with little continuity across various spatial and temporal scales (EEA 2020a; Hermoso et al. 2022; Santana et al. 2023; Geijzendorffer et al. 2016), and insufficient access to existing data makes it difficult for policymakers to make informed decisions and effectively design, implement and evaluate policies. A key first step to begin addressing these challenges and limitations is to provide a comprehensive assessment of current monitoring efforts in Europe, identify and address data gaps, workflow bottlenecks, and other challenges.

Many studies have focused on assessing these data needs and challenges with a data-focused approach, e.g., using remote sensing data or other (semi-)automated data collection methods (Proença et al. 2017; Vihervaara et al. 2017; Luque et al. 2018). Over-emphasizing such data-focused approaches runs the risk of neglecting the crucial socioeconomic and cultural contexts that motivate biodiversity monitoring (Kühl et al. 2020). Indeed, there is a lack of studies engaging directly with data users and policymakers to map their needs on biodiversity monitoring in Europe (e.g., through interviews, surveys, and meetings). After all, policies and data workflows are created by people, so there is a strong need to better understand the needs of data users and policymakers, and to identify relevant policy questions that rely on biodiversity monitoring. Additionally, due to the complexity of European biodiversity policy and the actors involved in data curation and analysis, there is a lack of comprehensive overviews of the European biodiversity monitoring landscape (Wetzel et al. 2018). In this paper, we employed with EuropaBON (https://europabon.org/) a user-centric four-step multi-stakeholder engagement process working with over 300 stakeholders across Europe to assess the current status and identify relevant policy questions, challenges, and possible solutions around biodiversity monitoring in Europe. This participatory approach closes a science-policy gap and brings data providers and end-users closer together. Based on suggestions by stakeholders, we propose five ways forward to addressing existing challenges and recommend the establishment of a European Biodiversity Observation Network through a permanent Biodiversity Monitoring and Coordinating Centre to improve the reporting and uptake of policy-relevant biodiversity data across Europe.

## 2. Methods

We conducted a four-step stakeholder engagement process to gather expert knowledge on biodiversity monitoring with over 300 stakeholders from the science and policy sector across Europe (see Figure 1). From the policy sector, experts from 18 EU member states with national contact points of the European Environment Information and Observation Network (Eionet) and eight European services, including experts from four Directorate-Generals of the European Commission, the European Environment Agency and Biodiversa+ took part. From society and science, experts from major natural history societies, such as the European Bird Census Council (EBCC), museums and data infrastructures as well as from universities and other research organisations participated. The stakeholder engagement process included (i) a public online stakeholder conference in May 2021, (ii) an open standardised online survey distributed across Europe (Moersberger et al. 2023a, 2023b) as well as (iii) targeted semi-structured interviews and (iv) an expert workshop in Sep 2021 with policy stakeholders.

**Figure 1.**
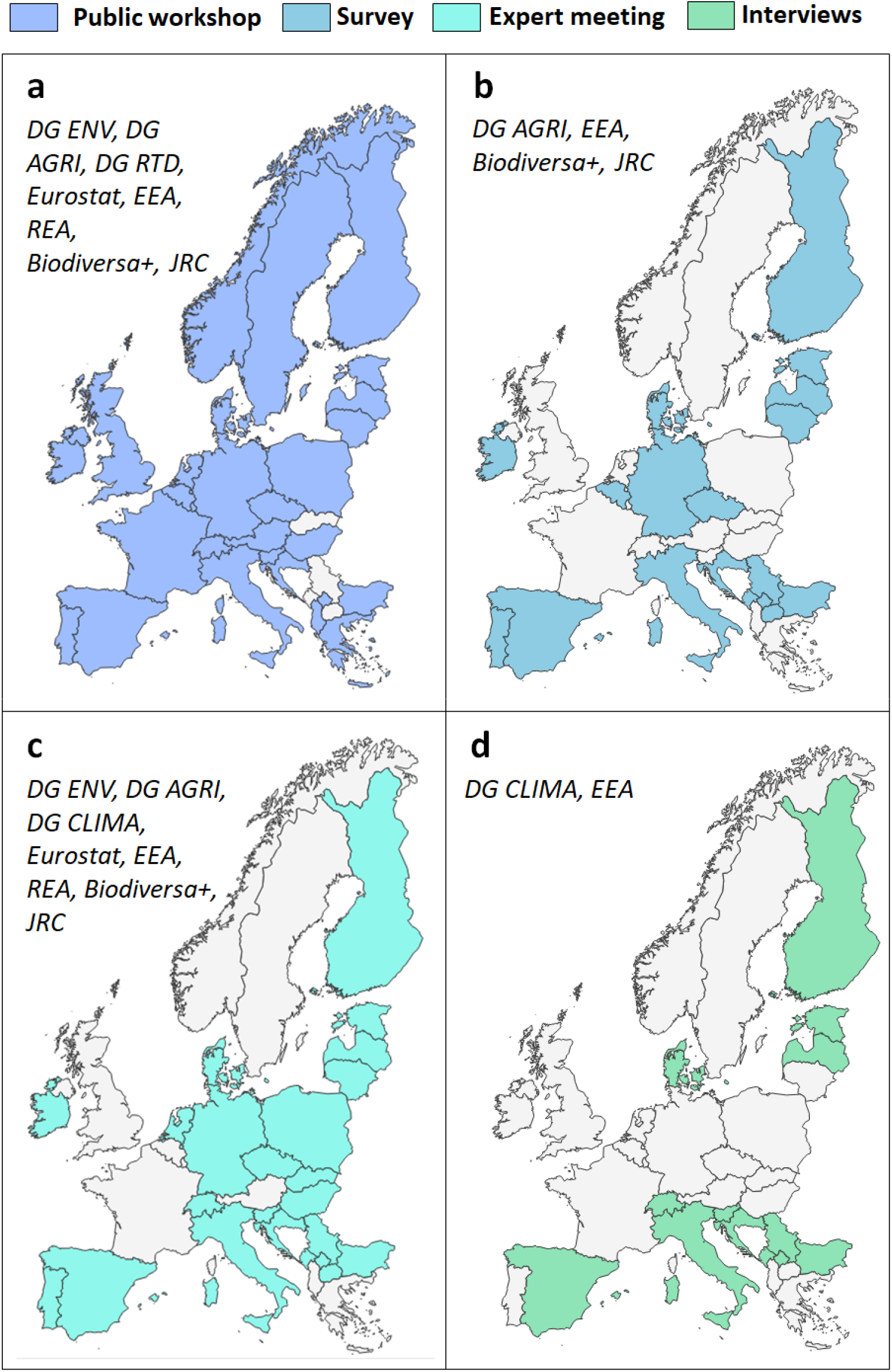
Geographic distribution of participants and European Union services present at the a) public stakeholder conference, b) surveys, c) expert meeting, and d) semi-structured interviews. European Commission services abbreviations: DG ENV (Directorate-General for Environment); DG AGRI (Directorate-General for Agriculture and Rural Development); DG CLIMA (Directorate-General for Climate Action); DG RTD (Directorate-General for Research and Innovation); EEA (European Environment Agency); REA (European Research Executive Agency); Biodiversa+ (European Biodiversity Partnership); JRC (Joint Research Centre).

Responses were coded with qualitative content analysis tool NVivo (Version 12; Edwards-Jones 2014; for details see S1). As with all coding studies, the identification of data groupings, codes, and themes remains subjective and open to different interpretations (Collier et al. 2012). The sample sizes, defined by the number of respondents willing to fill out the survey and participants in the workshops varied across the four methodological steps and bears the risk of self-selection bias. Freshwater and marine stakeholders were underrepresented, which renders the results for those realms less certain than those for terrestrial habitats. Additionally, the regional differences presented here should be viewed as a first step in the analysis of monitoring challenges in Europe. The four regions were represented by varying numbers of countries due to the unequal number of survey responses we received, which may induce some bias. Here, we focused mainly on national authorities and EU agencies, and further work is needed to assess the biodiversity monitoring needs of e.g. the private sector.

## 3. Results

### 3.1 The status quo: current national monitoring schemes in Europe

Our survey identified 274 biodiversity monitoring programmes that are currently conducted by European countries and agencies and shows a bias in biomes and taxonomic groups. Most schemes targeted terrestrial biodiversity (66%), followed by freshwater (24%) and marine/coastal biodiversity (21%). From a taxonomic perspective, birds were the most frequently monitored group, accounting for 28% of total monitoring efforts, primarily in support of the Birds Directive. Other groups with significant monitoring included mammals (18%) (particularly bats with 8%), insects (8%), plants (8%), pollinators (6%), and fish (6%). Representation of other taxonomic groups was low, ranging from 1% to 4%, with very little monitoring of microscopic taxa such as soil biota, fungi, plankton, and microorganisms (Figure 2). Genetic monitoring was mentioned as a largely underrepresented technique to inform on the state and trends of realms and taxa, with exceptions in countries such as Sweden and Switzerland, or on a local scale in Scotland.

**Figure 2.**
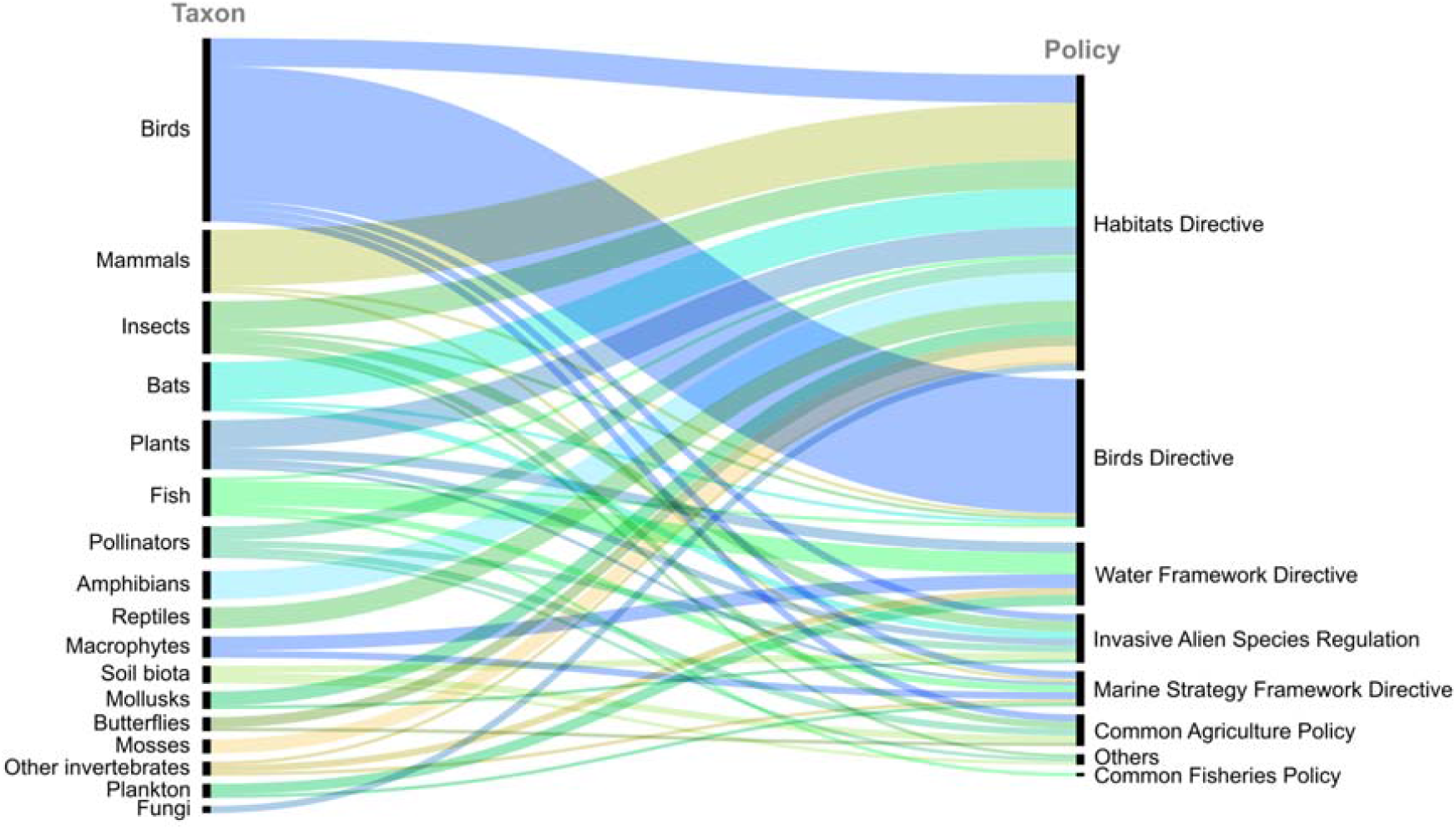
Data flows from monitoring taxonomic groups to informing various EU policies or directives. Each line in this figure represents one monitoring scheme reported in the survey from a total of N=274. Hence, the thickness of streams represents the number of monitoring schemes for a given taxonomic group. Taxonomic groups are shown here as listed by respondents, hence some smaller groupings may be included in larger groupings identified by the users during the assessment process. Taxonomic groups and policies are arranged in descending order of monitoring effort and reporting frequency, respectively.

With regards to reporting scale to policy directives the majority of schemes (62%) reported to the European level, to the national level (58%), and some to international conventions (16%) such as the Regional Seas Conventions, the Ramsar Convention, or the IUCN red list. The majority of respondents indicated that the EU or national policy biodiversity monitoring data are mainly used to report to the Habitats Directive (46%) and the Birds Directive (27%). Only a small portion of the data generated is used for other policies and directives such as the Common Agriculture Policy (CAP), Common Fisheries Policy, Water Framework Directive (WFD), Marine Strategy Framework Directive, or Invasive Alien Species Regulation (Figure 2).

### 3.2 Current policy uptake of biodiversity data

The most prominent use of biodiversity data in policy was related to the Habitats Directive and the Birds Directive with national applications for Species policies and management (55%), particularly for Species action plans (18%), Species management plans (13%), and Species conservation status (17%). Habitat conservation policies and management accounted for 31%, with Protected area management as the largest subcategory (10%). Biodiversity data were only moderately used for Land use management (11%), with Forest management plans and Land-use management plans as the largest subcategories (3% each). Meanwhile, cross-cutting topics such as Climate change and Ecosystem services were listed least often (3%) (Figure 3).

**Figure 3.**
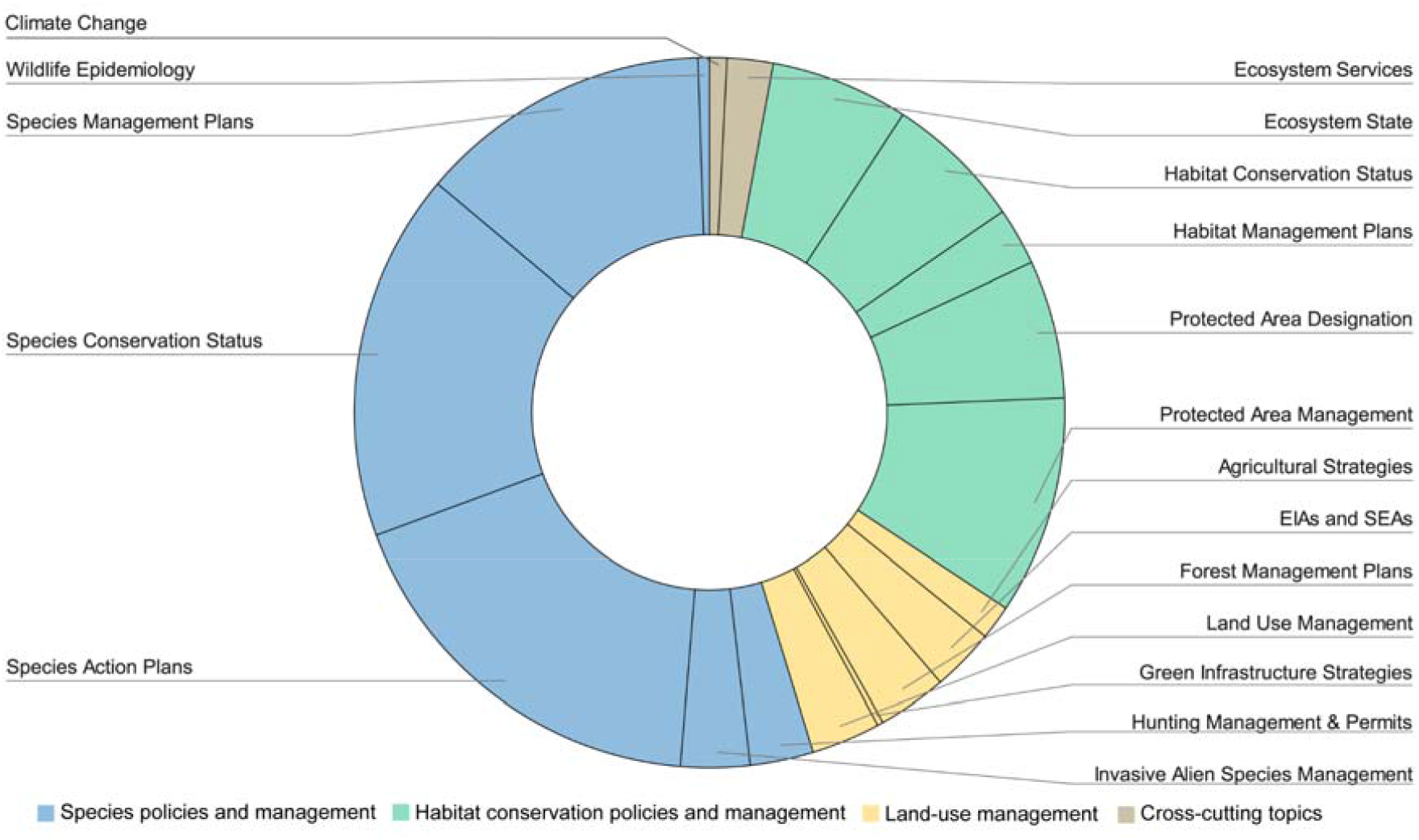
Policy uptake of data collected by biodiversity monitoring schemes in Europe, where each data segment represents the purpose of the respective biodiversity monitoring scheme listed by European countries and EU services.

Practically, our survey highlighted the regular uptake of monitoring data in national policy workflows. For example, in Estonia, biodiversity monitoring informs the regulation of hunting permits for wolves. Likewise, biodiversity monitoring has informed the designation of six new marine bird protection areas in Denmark. Biodiversity data are also gaining importance in the renewable energy sector, e.g. where bird monitoring, originally used for military aviation, is now being applied to wind energy to avoid collisions between migratory birds and wind turbines (Moersberger et al. 2022).

### 3.3 Key policy needs for future monitoring

The stakeholder conference participants identified four clusters of key policy questions related to biodiversity monitoring within the next decade (Table 1, for greater details, see Supplementary Table S2). The first cluster “Assessing biodiversity and species trends” focuses on understanding biodiversity status and trends, indicators for the quality of habitats, and assessing the impact of invasive species on the environment. These analyses are also needed to inform the second cluster “Biodiversity Policy Impact and Effectiveness” to assess the effectiveness of major biodiversity policies in Europe and the outcomes of conservation management and restoration. The third cluster “Integrating Biodiversity in other PolicySectors” branches out and focuses on the intersection of biodiversity conservation and other policy arenas such as agriculture, water management, climate change, green and blue infrastructure projects, poverty, equity, and trade. Finally, the fourth cluster “Operationalisation of Monitoring” explores ways to standardise and harmonise biodiversity monitoring programmes and integrate novel technologies to meet policy targets (Table 1). For more details on the policy questions and clusters, see Moersberger et al (2022).

**Table 1.** Four main clusters and thematic subcategories of selected policy questions regarding biodiversity monitoring within the next five to ten years as identified by European stakeholders.

| Topics | Selected Policy Questions |
| --- | --- |
| **I - Assessing Biodiversity and Species Trends** |  |
| **Understanding Biodiversity Trends** | How can we better assess biodiversity and species trends, and how does this affect ecosystem services? |
| **Indicators and Metrics** | How do we develop and measure indicators for habitat quality? |
|  | How can we produce reliable data-based risk- and impact assessments for habitat quality? |
| **Invasive species** | What are the impacts of invasive species on the environment and how can we better regulate them? |
| **II - Biodiversity Policy Impact and Effectiveness** |  |
| **Policy and Finance** | What is the effectiveness of major EU budgets and biodiversity policies in Europe, including support to the Natura 2000 network, species protection under the Habitats and Birds Directives, the WFD and the EU Biodiversity Strategy? |
|  | How can we 'future proof' biodiversity policy and identify harmful subsidies? |
| **Restoration** | How can we better assess where and how to restore biodiversity in Europe, with improved outcomes for the economy and society? |
|  | How can we monitor and restore terrestrial biodiversity and ecosystems outside areas that are subject to the Habitats Directive (mainly farmland, forest and urban areas)? |
| **Ecosystem Services** | How can we preserve biodiversity to maintain and use ecosystem services in a sustainable way? |
| **Marine Biodiversity** | How can we develop effective policies for marine biodiversity, which is susceptible to different patterns than terrestrial biodiversity? |
|  | How can major marine policies help track progress towards biodiversity targets, e.g. via the Common Fisheries Policy, Marine Strategy Framework Directive, EU Maritime Spatial Planning Directive, or the “new approach for a sustainable blue economy in the EU”? |
| **III - Integrating Biodiversity in other Policy Sectors** |  |
| **Agriculture and Farming** | How well is the Common Agricultural Policy (CAP) conserving/restoring biodiversity and how can agri-environment schemes be improved to enhance positive effects on biodiversity? |
|  | How does the Farm to Fork Strategy contribute to biodiversity, e.g., through pesticide reduction or organic farming? |
| **Water management** | How can we achieve 25,000 km of free-flowing rivers by 2030? |
|  | How to best distribute water allocations between agricultural and natural areas during drought, weighing economic benefits, vulnerabilities, and sustainable use? |
| **Climate Change Adaptation and Mitigation** | What is needed to take climate change impacts on biodiversity and ecosystem restoration into account? |
|  | What are the costs and benefits of policy targets toward climate change mitigation for biodiversity? |
| **Critical Green and Blue Infrastructure** | What is the effect of infrastructure projects on biodiversity (e.g., roads, wind farms, hydropower, power lines) and how can we work towards better green infrastructure? |
| **Telecoupling of negative effects** | How are European societies exporting negative externalities outside of Europe (e.g. deforestation) and how is this impacting biodiversity in Europe? |
| **Human health & Wellbeing** | How can we operationalise access to nature as a basic necessity? |
| **IV - Operationalisation of Monitoring** |  |
| **Financing** | How can we make monitoring of species and habitats economically viable, and how can we match the scale of funding with the urgency of biodiversity hotspot preservation? |
| **Data Standardisation and Harmonisation** | How can we seamlessly integrate and standardise monitoring, data flows, data products and policy across realms (marine, freshwater, terrestrial) and across the EU? |
|  | How can we better integrate underrepresented groups (e.g., plants, algae, invertebrates, soil organisms) in biodiversity monitoring? |
| **Digitisation and Novel Technologies** | How can we take advantage of novel technologies and digitalisation to meet biodiversity targets and support nature? |
| **New data sources** | How can we enhance (corporate) reporting (e.g., through Environmental Impact Assessments, Life Cycle Analyses) to improve biodiversity protection and restoration? |

### 3.4 Current monitoring challenges

The top ten challenges to current biodiversity monitoring in Europe, as listed in figure 4, were associated with four main types of obstacles: lack of integrated data, insufficient data, insufficient resources, and biased data (Figure 4). While lack of financial resources was ranked as the most important cross-cutting challenge by respondents across Europe, other challenges were ranked differently across different European regions. While Southern European countries emphasised the issue of insufficient data, such as limited spatial coverage, low monitoring frequency, and unavailability of raw data, Western European countries particularly highlighted the lack of long-term monitoring policies as well as the lack of human and technical capacities as their main challenges (Figure 4).

**Figure 4.**
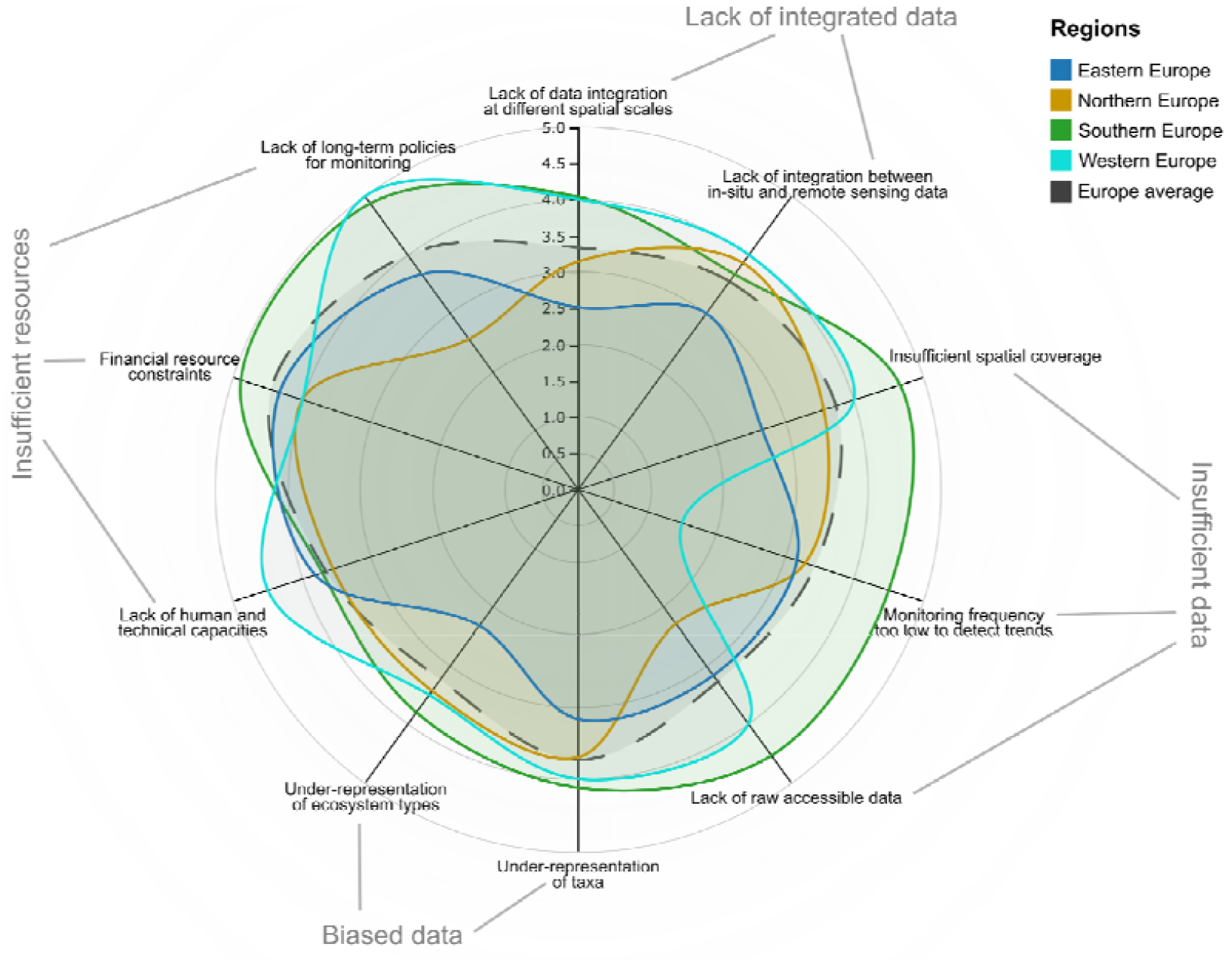
The ten most important challenges to biodiversity monitoring differ across the four European regions (information derived from surveys and interviews). Importance is ranked on a scale of 1 (least important) to 5 (most important). Importance ranks are averaged across countries in the four regions. Regional groupings are based on the geographic regions of the UN Statistics Division (United Nations 2019: Eastern Europe = Bulgaria, Hungary, Slovakia, Czechia, Poland. Southern Europe = Croatia, Montenegro, Serbia, Slovenia, Spain, Portugal, Kosovo, North Macedonia, Italy. Northern Europe = Denmark, Estonia, Finland, Latvia, Lithuania, Ireland. Western Europe = Switzerland, Netherlands, Germany.

### 3.5 Proposed avenues forward

To overcome these challenges in biodiversity monitoring, stakeholders proposed five potential ways forward: (i) enhanced coordination and cooperation, (ii) standardisation, enhanced data gathering and sharing, (iii) modelling and novel technologies, (iv) financial resources, and (v) capacity building and stakeholder engagement (Figure 5). It is important to note that financial resources are critical for the implementation of all ways forward, and long-term, targeted investments may be necessary for their successful realisation. The implementation of these potential solutions can also address the future key policy questions identified in section 3.2. Some proposed ways forward address several challenges simultaneously, and many are aligned with previous studies (Pe’er et al. 2021; Schmidt & Van der Sluis 2021). This emphasises their importance and the urgency of the transition toward implementation. The Europa Biodiversity Observation Network (EuropaBON) project is currently leading efforts to explore the feasibility and implementation of the suggested solutions in a Europe-wide coordinated approach (see also Moersberger et al. 2022).

**Figure 5.**
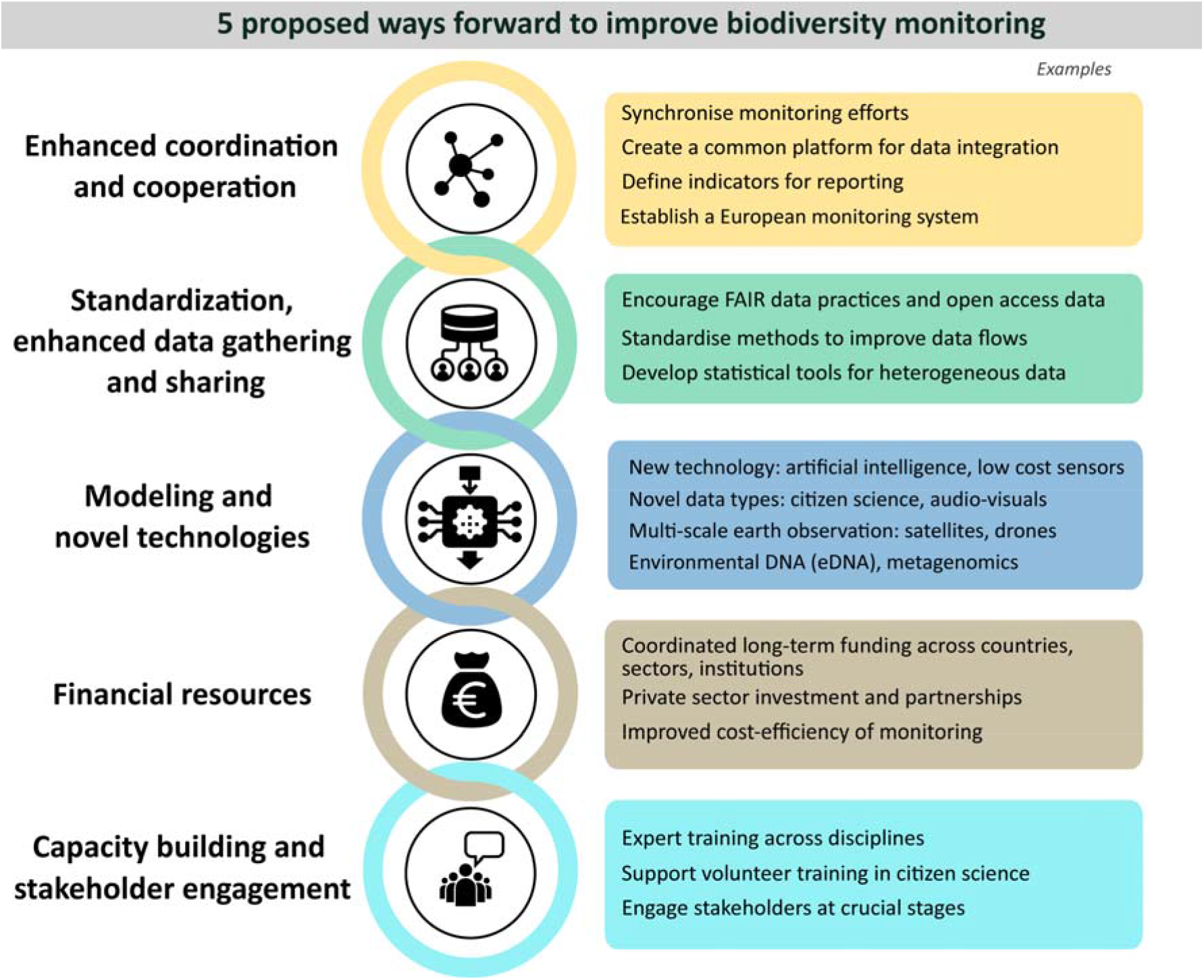
Five ways forward suggested by stakeholders to improve biodiversity monitoring and thereby policy impact in Europe.

## 4. Discussion

Our user and policy needs assessment reveals a rich, yet fragmented landscape of monitoring schemes in Europe, with biases towards certain biomes and taxonomic groups. The majority of national monitoring reports to the EU level with a predominant focus on the Habitats and Birds Directive with conservation management as the primary use for collected data, while transfer to other policy areas are yet underutilised. Key user and policy needs relate to assessing biodiversity trends, assessing biodiversity policy effectiveness, integrating biodiversity within other policy sectors and standardising monitoring programs. The top challenges to current biodiversity monitoring in Europe include inadequate data integration, biased data coverage, insufficient data availability and limited financial resources. To overcome these challenges, stakeholders suggested improving coordination and collaboration, standardising data collection and sharing, employing novel technologies, increasing financial resources, and enhancing capacity building and stakeholder engagement.

Our user-focused participatory approach through a four-step multi-stakeholder engagement served as a valuable tool for assessing current challenges and user needs of biodiversity monitoring and for developing user-and policy-relevant solutions to monitoring. The multi-stakeholder engagement process highlighted the need for comprehensive monitoring programmes that cover a wide range of biomes and taxonomic groups, beyond the dominant focus on terrestrial ecosystems and birds. This is in line with recent research that recognises monitoring biases towards charismatic species (Henle et al. 2013; Pilotto et al. 2020), easily surveyed species (Hermoso et al. 2017), and species listed in the Habitats and Birds Directives (Hoek 2022). Consequently, we currently have a limited understanding of the state and trends of biodiversity in Europe, insufficient monitoring evidence at the right resolution and temporal scales, and thus a gap between the information needs of policymakers and available data. However, stakeholders have clearly expressed their willingness to engage in more integrated and comprehensive approaches to biodiversity monitoring and have recognised the need for high-quality data, methods, and indicators to support effective conservation and restoration policy and, in the long term, ensure the sustainable management of different human activities and economic sectors.

The quantity and quality of baseline biodiversity datasets generated through current monitoring schemes differ across countries. Some countries do not have the capacity to conduct nationally-organised, government-funded biodiversity monitoring, while others monitor biodiversity, ecosystem state and processes through multiple schemes. Despite often mandatory reporting obligations, our survey and interview responses highlight that many countries currently struggle to fulfil the monitoring and reporting obligations of most European environmental directives. This is partly due to limited expertise, financial and human resources, as well as sub-nationally and locally differing schemes, taxonomies and habitat classifications. In line with these struggles, many countries expressed the wish to learn from their neighbours on how to enhance national biodiversity monitoring, e.g., through expert exchange groups and knowledge transfer platforms.

Additionally, we found that most of the biodiversity assessments performed by Member States under EU legislation are reported to the Habitats and Birds Directives, and their necessary re-use in other EU policies such as the Common Agriculture Policy, Common Fisheries Policy, Marine Strategy Framework Directive, Water Framework Directive, or Invasive Alien Species Regulation are limited or uncertain. This points to a potential data flow bottleneck, which could be addressed by improved data sharing policies and practices.

Despite these data flow limitations, our surveys have also drawn attention to numerous cases of successful biodiversity data uptake in policy workflows. We could assert that many monitoring schemes are crucial for informing species and habitat action plans, management plans, and conservation status appraisals, including some country- and context-specific insights such as use of bird data for wind energy planning.

Alongside success stories, a variety of challenges remain. The reporting for Article 17 of the Habitats Directive requires EU Member States to submit a report on the conservation status of habitat types and species to the European Commission every six years. Most countries regularly signal challenges related to poor data quality and completeness when drafting these reports. Currently, more than 60% of countries struggle with high levels of missing or unknown information and the methods for monitoring and reporting vary widely in rigour and precision (Ellwanger et al. 2018). Here, 11% of all Member States base more than half of their reporting on expert opinion or do not have any information on the methods used. The majority (82%) need to rely on combinations of methods, and only 7% state that more than half of their methods are actually based on complete surveys (EEA 2020). This lack of available data (or access to it) and performance reliance on sub-optimal methods implies that reporting results have a high level of uncertainty and are not very robust. The use of different methods also ensures that some data are not properly harmonised to derive EU assessments and indicators.

Our study results mirror the above mentioned challenges in current national and European monitoring efforts. The lack of detailed geo-referenced biodiversity data is severely hampering assessments of biodiversity and ecosystem trends, as well as infrastructure planning. Often, agencies only have access to regionally aggregated assessments of species or ecosystems and the underlying specific (raw) data are not traceable or easily accessible. Access to data from sectors such as agriculture, fisheries, and energy is also often limited, while at the same time valuable biodiversity information collected by those operators is not available, even for managers. Other challenges include a lack of guidance in identifying monitoring priorities; lack of standardised monitoring protocols; reluctance to change existing monitoring practices; and limited in-house knowledge and technical infrastructure to adequately mobilise and access biodiversity data (see also Mairota et al. 2015; Schmeller et al. 2015; Kühl et al. 2020). Although most countries responded that they use biodiversity monitoring data for at least some modelling to analyse future trends and (in a few cases) to support policy, a lack of capacity, expertise and funding currently prevents some countries from exploiting the full potential of biodiversity modelling. This indicates the need for more extensive capacity-building measures and expert knowledge exchange, which is one of the five recommendations for action proposed by stakeholders.

The effectiveness of these five ways forward will strongly depend on how they are implemented. Further research and actual tests are needed to explore the feasibility and potential impact of these solutions. In addition, it is pivotal to find effective ways of engaging stakeholders in their implementation and policy uptake. For example, the increased use of novel technologies could enhance cost-efficiency of data gathering and analysis. However, if the obligation to use novel technologies is imposed on agencies and individuals without the appropriate financial and capacity building measures, it can entail resistance to change. Several countries, especially those with smaller monitoring budgets and capacities, mentioned their fear of being overwhelmed by the obligation to use new technologies or methodologies due to lack of expertise. The proposed common platform for data integration (see Fig. 5) must be co-created with relevant stakeholders and embedded in existing platforms, e.g., the EBV data portal (iDiv 2023) and the Global Biodiversity Information Facility GBIF. Adopting the concept of Essential Biodiversity and Essential Ecosystem Services Variables (EBVs and EESVs) as standard monitoring variables can also address issues of data integration and scarcity, when accompanied by appropriate capacity building measures (see also Navarro et al. 2017).

Ultimately, the successful implementation of these potential solutions will require coordinated efforts across Europe. Our study aligns with previous research highlighting the need for a comprehensive assessment of current monitoring efforts and the creation of a centralised and integrated system to support biodiversity monitoring. We therefore recommend the establishment of a cost-efficient, user-oriented, policy-relevant, harmonised, and scalable European Biodiversity Observation Network. Such a network would help optimise existing observation efforts, harmonise data, provide financial and technical support to stakeholders across the continent, and improve our ability to predict and respond to key issues related to biodiversity loss. Its common framework should be carefully co-designed with all relevant users and policymakers, and it should be centrally coordinated.

The Europa Biodiversity Observation Network EuropaBON is currently working with a coalition of key stakeholders and a wide network of experts across Europe from science, agencies and natural history societies to establish such a network and to set up a permanent Biodiversity Monitoring Coordinating Centre in Europe. This centre can play a crucial role in facilitating cooperation, data mobilisation and integration, data interoperability, the alignment and design of new sampling methods, the provision of workflows to generate standardised biodiversity indicators, and the provision of capacity building. Hence, the Biodiversity Monitoring Coordinating Centre would serve as a powerful tool to facilitate and boost the five avenues forward identified in this study. With the expected forthcoming EU Restoration Law, these needs are amplified. By effectively coordinating efforts across countries, relevant European agencies, and non-governmental organisations, the centre could help foster and drive effective, collaborative biodiversity monitoring in Europe. Integrative monitoring will be key to effectively inform and respond to urgent 2030 policy goals for sustainable biodiversity management,conservation and restoration. Effective biodiversity monitoring in Europe will be critical in addressing the future key policy questions identified in this study and to provide accountability for policy effectiveness and impact within and across policy sectors.

## Acknowledgements

We are indebted to all participants in our surveys, interviews and workshops for their deep insights into monitoring needs, obstacles and possible solutions. We also thank Maria Dornelas for her advice and contributions to EuropaBON. The EuropaBON project receives funding from the European Union’s Horizon 2020 research and innovation programme under grant agreement No 101003553. We gratefully acknowledge the support of the German Centre for Integrative Biodiversity Research (iDiv) funded by the German Research Foundation (DFG-FZT 118, 202548816). CC acknowledges support from the Portuguese Foundation for Science and Technology through funding to CEG/IGOT Research Unit (UIDB/00295/2020 and UIDP/00295/2020). The views and opinions expressed in this article are those of the authors and do not necessarily reflect the official position of the European Commission.

## References

Collier, D., LaPorte, J. & Seawright, J. (2012). Putting Typologies to Work: Concept Formation, Measurement, and Analytic Rigor. Polit. Res. Q., 65, 217–232.

COP to the CBD. (2022). Kunming-Montreal Global biodiversity framework, Draft decision submitted by the President. Montreal, Canada.

Directorate-General for Environmen. (2021). EU biodiversity strategy for 2030: bringing nature back into our lives. Publications Office of the European Union, LU.

Directorate-General for Environment. (2022). Proposal for a Nature Restoration Lazw.

Edwards-Jones, A. (2014). Qualitative data analysis with NVIVO.

EEA. (2020). Data quality & completeness for Article 17 [WWW Document]. URL https://tableau.discomap.eea.europa.eu/t/Natureonline/views/DataqualitycompletenessforArticle17/Article17Dataqualitycompleteness?:showAppBanner=false&:display_count=n&:showVizHome=n&:origin=viz_share_link&:isGuestRedirectFromVizportal=y&:embed=y

Ellwanger, G., Runge, S., Wagner, M., Ackermann, W., Neukirchen, M., Frederking, W., Müller, C., Ssymank, A. & Sukopp, U. (2018). Current status of habitat monitoring in the European Union according to Article 17 of the Habitats Directive, with an emphasis on habitat structure and functions and on Germany. Nat. Conserv., 29, 57–78.

Geijzendorffer, I.R., Regan, E.C., Pereira, H.M., Brotons, L., Brummitt, N., Gavish, Y., Haase, P., Martin, C.S., Mihoub, J.-B., Secades, C., Schmeller, D.S., Stoll, S., Wetzel, F.T. & Walters, M. (2016). Bridging the gap between biodiversity data and policy reporting needs: An Essential Biodiversity Variables perspective. J. Appl. Ecol., 53, 1341–1350.

Henle, K., Bauch, B., Auliya, M., Külvik, M., Pe’er, G., Schmeller, D.S. & Framstad, E. (2013). Priorities for biodiversity monitoring in Europe: A review of supranational policies and a novel scheme for integrative prioritization. Ecol. Indic., Biodiversity Monitoring, 33, 5–18.

Hermoso, V., Clavero, M., Villero, D. & Brotons, L. (2017). EU’s Conservation Efforts Need More Strategic Investment to Meet Continental Commitments. Conserv. Lett., 10, 231–237.

Hoek, N. (2022). A Critical Analysis of the Proposed EU Regulation on Nature Restoration: Have the Problems Been Resolved? Eur. Energy Environ. Law Rev., 31.

iDiv. (2023). EBV Data Portal [WWW Document]. URL https://portal.geobon.org/home

Kühl, H.S., Bowler, D.E., Bösch, L., Bruelheide, H., Dauber, J., Eichenberg, D., Eisenhauer, N., Fernández, N., Guerra, C.A., Henle, K., Herbinger, I., Isaac, N.J.B., Jansen, F., König-Ries, B., Kühn, I., Nilsen, E.B., Pe’er, G., Richter, A., Schulte, R., Settele, J., van Dam, N.M., Voigt, M., Wägele, W.J., Wirth, C. & Bonn, A. (2020). Effective Biodiversity Monitoring Needs a Culture of Integration. One Earth, 3, 462–474.

Luque, S., Pettorelli, N., Vihervaara, P. & Wegmann, M. (2018). Improving biodiversity monitoring using satellite remote sensing to provide solutions towards the 2020 conservation targets. Methods Ecol. Evol., 9, 1784–1786.

Mairota, P., Cafarelli, B., Didham, R.K., Lovergine, F.P., Lucas, R.M., Nagendra, H., Rocchini, D. & Tarantino, C. (2015). Challenges and opportunities in harnessing satellite remote-sensing for biodiversity monitoring. Ecol. Inform., 30, 207–214.

Moersberger, H., Martin, J.G., Junker, J., Georgieva, I., Bauer, S., Beja, P., Breeze, T., Brotons, L., Bruelheide, H., Fernández, N., Fernandez, M., Jandt, U., Langer, C., Lyche Solheim, A., Maes, J., Moreira, F., Pe’er, G., Santana, J., Shamoun-Baranes, J., Smets, B., Valdez, J., McCallum, I., Pereira, H.M. & Bonn, A. (2022). Europa Biodiversity Observation Network: User and Policy Needs Assessment. ARPHA Prepr., 3, e84517.

Moersberger, H., Martin, J.G.C., Junker, J., Georgieva, I., Maes, J., McCallum, I., Pereira, H.M. & Bonn, A. (2023a). National survey to co-design the Europa Biodiversity Observation Network (EuropaBON). ARPHA Prepr., 4, ARPHA Preprints.

Moersberger, H., Martin, J.G.C., Junker, J., Georgieva, I., Maes, J., McCallum, I., Pereira, H.M. & Bonn, A. (2023b). European survey to co-design the Europa Biodiversity Observation Network (EuropaBON). ARPHA Prepr., 4, ARPHA Preprints.

Navarro, L.M., Fernández, N., Guerra, C., Guralnick, R., Kissling, W.D., Londoño, M.C., Muller-Karger, F., Turak, E., Balvanera, P. & Costello, M.J. (2017). Monitoring biodiversity change through effective global coordination. Curr. Opin. Environ. Sustain., 29, 158–169.

Pe’er, G., Birkenstock, M., Lakner, S. & Röder, N. (2021). The Common Agricultural Policy post-2020: Views and recommendations from scientists to improve performance for biodiversityr1: Volume 3, Policy Brief.

Perino, A., Pereira, H.M., Felipe-Lucia, M., Kim, H., Kühl, H.S., Marselle, M.R., Meya, J.N., Meyer, C., Navarro, L.M., van Klink, R., Albert, G., Barratt, C.D., Bruelheide, H., Cao, Y., Chamoin, A., Darbi, M., Dornelas, M., Eisenhauer, N., Essl, F., Farwig, N., Förster, J., Freyhof, J., Geschke, J., Gottschall, F., Guerra, C., Haase, P., Hickler, T., Jacob, U., Kastner, T., Korell, L., Kühn, I., Lehmann, G.U.C., Lenzner, B., Marques, A., Motivans Švara, E., Quintero, L.C., Pacheco, A., Popp, A., Rouet-Leduc, J., Schnabel, F., Siebert, J., Staude, I.R., Trogisch, S., Švara, V., Svenning, J.-C., Pe’er, G., Raab, K., Rakosy, D., Vandewalle, M., Werner, A.S., Wirth, C., Xu, H., Yu, D., Zinngrebe, Y. & Bonn, A. (2022). Biodiversity post-2020: Closing the gap between global targets and national-level implementation. Conserv. Lett., 15, e12848.

Pilotto, F., Kühn, I., Adrian, R., Alber, R., Alignier, A., Andrews, C., Bäck, J., Barbaro, L., Beaumont, D., Beenaerts, N., Benham, S., Boukal, D.S., Bretagnolle, V., Camatti, E., Canullo, R., Cardoso, P.G., Ens, B.J., Everaert, G., Evtimova, V., Feuchtmayr, H., García-González, R., Gómez García, D., Grandin, U., Gutowski, J.M., Hadar, L., Halada, L., Halassy, M., Hummel, H., Huttunen, K.-L., Jaroszewicz, B., Jensen, T.C., Kalivoda, H., Schmidt, I.K., Kröncke, I., Leinonen, R., Martinho, F., Meesenburg, H., Meyer, J., Minerbi, S., Monteith, D., Nikolov, B.P., Oro, D., Ozoliņš, D., Padedda, B.M., Pallett, D., Pansera, M., Pardal, M.Â., Petriccione, B., Pipan, T., Pöyry, J., Schäfer, S.M., Schaub, M., Schneider, S.C., Skuja, A., Soetaert, K., Spriņge, G., Stanchev, R., Stockan, J.A., Stoll, S., Sundqvist, L., Thimonier, A., Van Hoey, G., Van Ryckegem, G., Visser, M.E., Vorhauser, S. & Haase, P. (2020). Meta-analysis of multidecadal biodiversity trends in Europe. Nat. Commun., 11, 3486.

Proença, V., Martin, L.J., Pereira, H.M., Fernandez, M., McRae, L., Belnap, J., Böhm, M., Brummitt, N., García-Moreno, J., Gregory, R.D., Honrado, J.P., Jürgens, N., Opige, M., Schmeller, D.S., Tiago, P. & van Swaay, C.A.M. (2017). Global biodiversity monitoring: From data sources to Essential Biodiversity Variables. Biol. Conserv., 213, 256–263.

Santana, J., Porto, M., Brotons, L., Junker, J., Kissling, W.D., Lumbierres, M., Moe, J., Morán-Ordóñez, A., Pereira, H., Lyche Solheim, A., Villero, D., Moreira, F. & Beja, P. (2023). D3.2 Report on gaps and important new areas for monitoring in Europe. ARPHA Prepr., 4, ARPHA Preprints.

Schmeller, D.S., Julliard, R., Bellingham, P.J., Böhm, M., Brummitt, N., Chiarucci, A., Couvet, D., Elmendorf, S., Forsyth, D.M. & Moreno, J.G. (2015). Towards a global terrestrial species monitoring program. J. Nat. Conserv., 25, 51–57.

Schmidt, A.M. & Van der Sluis, T. (2021). Improving the Availability of Data and Information on Species, Habitats and Sites. European Commission, Directorate-General Environment, Wageningen, The Netherlands.

United Nations. (2019). Standard country or area codes for statistical use (M49). Statistical Services Branch, UN Statistics Division, New York.

Vihervaara, P., Auvinen, A.-P., Mononen, L., Törmä, M., Ahlroth, P., Anttila, S., Böttcher, K., Forsius, M., Heino, J., Heliölä, J., Koskelainen, M., Kuussaari, M., Meissner, K., Ojala, O., Tuominen, S., Viitasalo, M. & Virkkala, R. (2017). How Essential Biodiversity Variables and remote sensing can help national biodiversity monitoring. Glob. Ecol. Conserv., 10, 43–59.

Wetzel, F.T., Bingham, H.C., Groom, Q., Haase, P., Koljalg, U., Kuhlmann, M., Martin, C.S., Penev, L., Robertson, T., Saarenmaa, H., Schmeller, D.S., Stoll, S., Tonkin, J.D. & Haeuser, C.L. (2018). Unlocking biodiversity data: Prioritization and filling the gaps in biodiversity observation data in Europe. Biol. Conserv., 221, 78–85.

