## Supplementary for "Biodiversity monitoring in Europe: user and policy needs"

**Supporting information**

Supplement S1: Methodology on stakeholder engagement process

A three-day online public stakeholder workshop was held in May 2021, and attended by 246 participants from across Europe and elsewhere. Theparticipants were mainly from the academic sector, non-governmental organisations (NGOs), or the policy sector. During the workshop, participants identified key policy questions that require biodiversity monitoring over the next 5-10 years to inform the EU Biodiversity Strategy and other policies. Participants also listed and ranked current challenges and potential solutions, which were further evaluated in using the software EasyRetro (<https://easyretro.io/>) and in subsequent steps in the stakeholder engagement process.

We then  conducted a standardised online survey to identify biodiversity monitoring initiatives implemented by European countries and organisations, along with current challenges and potential solutions (Moersberger et al. 2023a, 2023b). The survey was sent to all national contact points of the European Environment Information and Observation Network (Eionet), nine EU services, and other relevant experts working at the interface of biodiversity monitoring and policy in 37 European countries. Respondents were asked to list and categorise monitoring schemes based on reporting scale, alignment with EU directives, biomes and taxonomic groups, and their main purpose (e.g., management strategies or designation of conservation areas). Policy and research experts from 23 European countries and four EU services responded between July and September 2021. A list of national and European monitoring schemes was extracted from the surveys and classified using the qualitative content analysis tool NVivo (Version 12; Edwards-Jones 2014). A grounded theory approach was used in the analyses (Birks & Mills 2022), meaning that where possible classifications (or ‘codes’) emerged from data inductively, allowing a more detailed grouping of statements (e.g. hunting management and permits), than would be possible by applying an existing framework to the data (Strauss & Corbin 1990). This method allows more granularity and tangibility from survey data, rather than over-generalizing classification results. Categories and data groupings of the purpose of data collection emerged from the data itself, meaning that the most detailed grouping possible was used when reported by respondents (e.g., hunting management and permits). When broader purposes were reported, these were used as such (e.g., species action plans), and where necessary, were merged based on guidelines from the United Nations Economic Commission for Europe (2023).

Third, to complement and validate survey results, semi-structured interviews were conducted from September to December 2021 with a sub-sample of those who responded to the survey (experts from 13 European countries and two EU services). Participation in the interviews was based on the availability and willingness of the respondents, following personal invitation and reminders. Insights from the interview transcriptions inform the discussion section of this paper.

As the fourth and last step, we held an online expert meeting with 40 policy experts from 18 European countries and eight EU services in September 2021. All Eionet national contact points and all other experts who participated in the survey were invited. The expert meeting served to gather feedback, validate the survey and interview analyses, and facilitate trustful, in-depth discussions among policy experts. EU services that participated were the European Research Executive Agency, DG AGRI, DG Environment, DG Climate, DG Research & Innovation, the European Environment Agency, the Biodiversa + Partnership and the Joint Research Center.

Supplement S2: Key policy questions as identified by stakeholders

Supplementary Table 2. Four main clusters and subcategories of key policy questions regarding biodiversity monitoring within the next five to ten years as identified by European stakeholders.

| **Cluster** | **Sub category** | **Policy question** |
| --- | --- | --- |
| Integrated cross-sectoral policies | General questions | How do we better integrate biodiversity policies with interconnected sectors - e.g. food, farming, diets, energy, water management, pollution, climate, poverty, equity? |
|  |  | How to balance the various policy needs and decisions (and hence subsidiaries) for the benefit of ecosystem health? |
|  |  | How can we 'future proof' biodiversity policy? |
|  | Common Agricultural Policy and agriculture | What are the key pollinators of different crops in the EU that should be integrated by the CAP? |
|  |  | How is biodiversity change driving changes in agricultural soils and how will this impact sustainable farming targets? |
|  |  | How well is CAP conserving/restoring biodiversity and how can agri-environment schemes be improved to enhance positive effects on biodiversity? |
|  |  | How does the Farm to Fork strategy contribute to biodiversity, e.g. through pesticide reduction, organic farming etc.? |
|  | Climate change | What is needed to take climate change impacts on biodiversity into account? |
|  |  | How can we restore ecosystems in a climate change perspective? |
|  |  | What are the costs and benefits of climate change mitigation policies targets for biodiversity, including eventual negative impacts (e.g., converting species-rich grasslands to forests for carbon sequestration)? |
|  |  | What is the climate dependency of goals/targets set by Nature Directives, WFD, and MSFD? |
|  |  | How can we improve the links between climate policies and biodiversity policies? |
|  | Infrastructure | What is the effect of infrastructure projects on biodiversity (e.g. roads, wind farms, power lines)? |
|  |  | How can we measure and work towards better green infrastructure (connectivity for species)? |
|  | Freshwater/Water Framework Directive | What is the best approach to distribute water allocations between agricultural and natural area's during droughts, weighting economic benefits, vulnerabilities and sustainable use? |
|  |  | How can we better link policy on Agriculture and WFD with Natura 2000? |
|  |  | How can we achieve 25,000km of free-flowing rivers by 2030? |
|  |  | What is the contribution of the Water Framework Directive to the conservation of biodiversity and ecosystem services? |
|  |  | How best to establish groundwater biodiversity assessment/monitoring schemes related to the WFD and Groundwater Directives? |
|  | Human-nature connections | How do we measure effects of forest practices on the status of forest biodiversity? |
|  |  | How can we operationalise access to nature as a basic necessity for people in the EU and the world? |
|  |  | How to manage and protect wildlife biodiversity despite human activities? |
| Policy impacts and effectiveness | Effectiveness and impact of policies and measures | What is the effectiveness of major biodiversity policies in Europe, incl. Natura 2000, species protection in the Habitats and Birds Directives, and the EU Biodiversity Strategy? |
|  |  | What is the effectiveness of EU budgets, e.g. what are the outcomes of expenditures on nature conservation and restoration? |
|  |  | How can we produce reliable data-based risk and impact assessments? |
|  |  | How can policy decisions better be linked to biodiversity indicators? |
|  |  | What is the most effective way to distribute government subsidies to ensure they deliver biodiversity outcomes (e.g. via CAP)? |
|  |  | How can corporate reporting (e.g. through EIA, LCA) improve biodiversity protection and restoration? |
|  | Conservation/  protection | How can we more effectively preserve, protect, and increase natural areas? |
|  |  | How do we ensure at least 30% of habitat and species are in favourable conservation status? |
|  | Restoration of biodiversity and ecosystems | How can we monitor and restore terrestrial biodiversity and ecosystems outside of the Habitats directive (mainly farmland, forest and urban areas)? |
|  |  | How can we better assess where and how to restore biodiversity in Europe? |
|  |  | How does biodiversity restoration action result in improved outcomes for the economy and society? |
|  |  | Is the money for monitoring and observation spent just to observe biodiversity loss or also on actions to halt and restore biodiversity? |
|  | Ecosystem services | How can we preserve biodiversity to maintain ecosystem services? |
|  |  | To what extent is insect diversity and biomass in agricultural, urban, and natural habitats declining (or recovering) and how does this affect ecosystem services (e.g. pollination, pest control, human well-being)? |
|  |  | How can we use ecosystems and their services in a sustainable way? |
|  | Telecoupling | How are European societies exporting negative externalities outside of Europe and how is this impacting biodiversity IN Europe? |
|  |  | How can we ensure that Europe’s policies do not undermine biodiversity elsewhere? |
|  |  | How can the EU mitigate the impacts of their trade with the rest of the world, namely regarding the source of raw materials from tropical and other biodiversity-rich countries? |
|  | Marine biodiversity | How to develop effective policies for marine biodiversity, which is susceptible to different patterns than terrestrial biodiversity, spanning entire countries' EEZ and also ABNJ? |
| Operationalisation of future policies and measures | Funding/Financing | How can we make land monitoring on species and habitats economically viable? |
|  |  | How can we better match funding with biodiversity hotspot preservation? |
|  |  | How can we identify “perverse” funding, e.g. through subsidies? |
|  | Digitalisation & novel technologies | How do we take advantage of novel technology and digitalisation to meet biodiversity targets and support nature? |
|  | Societal dynamics & engagement | How do we deal with a dynamic society that changes priorities every generation? |
|  |  | How to motivate a broader range of people to participate in bending the biodiversity curve by making them able to meaningfully contribute? |
| Knowledge, research, coordinated monitoring | Biodiversity trends/  Understanding biodiversity | How can we stop or reverse biodiversity loss, how can we address the main drivers? |
|  |  | Which is the impact of biodiversity on human beings and how can this be measured? |
|  |  | How can we better integrate underrepresented groups (e.g., invertebrates, soil organisms) in biodiversity monitoring? |
|  |  | How do we measure and create indicators for the quality of habitats? |
|  |  | How do we know if biodiversity and abundance of organisms are increasing (invasives) or decreasing (loss of species)? |
|  |  | What is the impact of invasive alien species on the environment? |
|  | Monitoring data & method integration | How do we successfully and seamlessly integrate monitoring, data flow, data products and policy across realms (marine, freshwater, terrestrial, aerial)? |
|  |  | How can biodiversity monitoring programmes be standardised in the EU? |
|  |  | Are birds and butterflies sufficient indicators? Or do we need to include ecosystem composition - covering a broader taxonomic group in order to create effective policy interventions. |

**References**

Birks, M. & Mills, J. (2022). *Grounded theory: A practical guide*. Sage.

Edwards-Jones, A. (2014). Qualitative data analysis with NVIVO.

Moersberger, H., Martin, J.G.C., Junker, J., Georgieva, I., Maes, J., McCallum, I., Pereira, H.M. & Bonn, A. (2023a). National survey to co-design the Europa Biodiversity Observation Network (EuropaBON). *ARPHA Prepr.*, 4, ARPHA Preprints.

Moersberger, H., Martin, J.G.C., Junker, J., Georgieva, I., Maes, J., McCallum, I., Pereira, H.M. & Bonn, A. (2023b). European survey to co-design the Europa Biodiversity Observation Network (EuropaBON). *ARPHA Prepr.*, 4, ARPHA Preprints.

Strauss, A. & Corbin, J.M. (1990). *Basics of qualitative research: Grounded theory procedures and techniques*. Basics of qualitative research: Grounded theory procedures and techniques. Sage Publications, Inc, Thousand Oaks, CA, US.

United Nations Economic Commission for Europe. (2023). *Guidelines for developing national biodiversity monitoring systems*. United Nations Publications, New York, USA.
